# Presurgical language fMRI: Current practices and patient outcomes in epilepsy surgical planning

**DOI:** 10.1101/185835

**Authors:** Christopher F.A. Benjamin, Alexa X Li, Hal Blumenfeld, R Todd Constable, Rafeed Alkawadri, Stephan Bickel, Christoph Helmstaedter, Stefano Meletti, Richard Bronen, Simon K. Warfield, Jurriaan M. Peters, David Reutens, Monika Połczyńska, Dennis D. Spencer, Lawrence J. Hirsch

**Affiliations:** Yale University School of Medicine, United States of America; Quinnipiac University School of Medicine, United States of America; Stanford University, United States of America; University of Bonn, Germany; University of Modena and Reggio Emilia, Italy; Harvard Medical School, United States of America; The University of Queensland, Australia; University of California Los Angeles, United States of America.

## Abstract

The goal of this study was to document current clinical practice and report patient outcomes in presurgical language functional MRI (fMRI) for epilepsy surgery. Epilepsy surgical programs worldwide were surveyed as to the utility, implementation, and efficacy of language fMRI in the clinic; 82 programs responded between July 2015 and January 2016. Respondents were predominantly from the US (61%), were academic programs (85%), and predominantly evaluated adults (44%), both adults and children (40%), or children only (16%). Nearly all (96%) respondents reported using language fMRI. fMRI is used for language lateralization (100%) and for localizing (44%) language cortex to guide surgical margins. While typically considered useful, programs often reported at least one instance of disagreement with other measures (56%). When used to localize language cortex, direct brain stimulation typically confirmed fMRI findings (74%) but instances of unpredicted decline were reported by 17% of programs and unexpected preservation of function were reported by 54%. Programs reporting unexpected decline did not clearly differ from those which did not. Clinicians using fMRI to guide surgical margins typically map Broca’s and Wernicke’s areas but not other known language areas. Language fMRI is widely used for lateralizing language cortex in patients with medically intractable epilepsy. It is also frequently used to localize language cortex though it is not yet well validated for this purpose. Many centers report cases of unexpected language preservation when fMRI activation is resected, and cases of language decline when it is not. Care will almost certainly be improved by standardizing protocols and accurately detailing the relationship between fMRI-positive areas and post-surgical language decline.

## 1.1 Introduction

Neurosurgery is an effective and potentially curative treatment for temporal lobe epilepsy.^1^ Surgical risk to language and memory can exclude a patient from treatment. As 34%-41% of left temporal patients undergoing focal resections experience a decline in naming,^2,3^ determining the surgical risk to language remains essential.

While the Intracarotid Amobarbital Test ("Wada" testing) has been the gold standard for determining the language dominant hemisphere, functional magnetic resonance imaging (fMRI) can be accurate^4^ and less costly,^5^ and is non-invasive. The evidence supporting fMRI’s validity was recently outlined,^6^ with the conclusion that language fMRI is a valid alternative to Wada testing in most patients. One approach to incorporating fMRI in clinical decision making developed by Swanson and colleagues.^7^ In short, if language fMRI shows left hemisphere dominance and a patient has right hemisphere pathology, Wada testing may be deemed unnecessary for language lateralization. Given the significance of a right hemisphere language finding and the increased probability of discordance with Wada,^8^ an argument can be made for repeating language fMRI for right dominance findings.

While different language fMRI tasks will yield differing maps and accuracy in predicting post-operative decline,^9^ good estimates of fMRI’s validity in a tertiary epilepsy setting are available for a semantic decision making task. With this protocol, a key study found decline of more than two standard deviations in naming skill (relative to controls) could be predicted with 100% sensitivity and 73% specificity, with prediction superior to that using Wada (92% and 45%, respectively).^4^ Overall, 41% of variance in post-operative language skill was predicted by fMRI and the positive predictive value was 81% for fMRI and 67% for Wada. In 229 epilepsy patients, using a slightly different analysis, 80% were classified as left dominant using fMRI, of which 92% were also Wada left dominant (167/182; 15 bilateral).^8^ fMRI bilateral cases (n=28) were typically left (46%) or bilateral (36%) on Wada, though occasionally (18%) right. fMRI right cases were most often also right on Wada (53%), though could be markedly discrepant (21% [n=4] left Wada; all right handed, right seizure foci). Explicitly examining cases of fMRI—Wada disagreement, this protocol has been shown to more accurately predict naming decline than Wada testing.^4^ BOLD fMRI language maps are not validated for the routine drawing of boundaries around indispensable cortex (the removal of which is associated with language decline), however.

For clinicians, guidelines for fMRI use are available within radiology^10^ and neuropsychology.^11^ Recent surveys from the European Union’s E-PILEPSY project showed 82% of European epilepsy programs use language fMRI,^12^ with Wada being used more judiciously.^12^ The questions clinical teams ask of language fMRI, how the results are interpreted, and whether they improve clinical care, however, remain unclear. Whether current clinical practices are in keeping with current best evidence is also unknown.

The goals of the current study were to characterize how epilepsy programs use language fMRI in surgical planning and aggregate reports of patient outcomes otherwise unlikely to be reported, to help inform future research and clinical fMRI guideline development.

## 2.1 Methods

This study was approved and overseen by Yale Medical School’s Institutional Review Board. All participants provided informed consent (Supplement 1).

## 2.2 Survey

A survey focused on clinicians’ use of and experience with presurgical language fMRI was developed (Supplement 1). Questions centered on identifying the dominant hemisphere ("lateralization"); identifying language areas to guide surgical margins ("localization"); how programs use fMRI; their confidence in results; and patient outcomes. A past survey of extraoperative mapping was used as a reference in design.^13^ Clinicians from neurology, radiology and neurosurgery provided feedback on clarity and length. The research design was reviewed by Yale’s Center for Analytic Sciences. A second, related survey of those acquiring fMRI data will be reported separately. The survey was presented via www.qualtrics.com. Questions were organized hierarchically with all respondents answering key questions which could elicit related questions. If a question was not answered a warning appeared, but the respondent could elect to continue without answering.

## 2.3 Procedure

### 2.3.1 Site identification

We invited all level 3 and 4 epilepsy centers in the US National Association of Epilepsy Centers (NAEC) and contacted epilepsy organizations (e.g., ILAE members) and prominent researchers worldwide. We asked individuals to invite other programs they knew, to increase sample size, using a modified "snowball sampling" approach.^14^

### 2.3.2 Data collection

The survey was open 07.17.2015-01.15.2016. We emailed invitations to all NAEC sites and followed up by telephone. We also contacted researchers, epilepsy organizations, and ILAE member committees worldwide requesting they forward an email invitation to relevant contacts. In 11.2015 we sent reminders and emailed the American Epilepsy Society (AES) listserv.

### 2.3.3 Final sample

82 surveys were received from respondents involved in selecting patients for epilepsy surgery. The number of responses per question is indicated in square brackets; e.g., [n=X]. Responses vary due to the study’s hierarchical design; some specific responses elicited additional relevant questions. Data were cleaned; of note, two respondents answered only initial questions on their confidence in methods for language lateralization, and three reported no patients received fMRI. Respondent’s country was estimated from stated location (per the survey or email) or IP address. Descriptive statistics are presented and compared using t and Fisher’s Exact tests.

Programs were typically university-affiliated (85%), evaluating a mean of 106 (SD 67; 10-300) patients annually and surgically treating a mean of 34 (SD 21, 0-100) [n=80 respondents]. Overall 84% evaluated adults and 56% children; specifically - 44% evaluated predominantly adults; 40% predominantly both adults and children; and 16% predominantly children (<18 years) [n=80]. Respondents who identified their background included neurologists (89%), neurosurgeons (6%) and neuropsychologists (5%) [n=66]. They were typically clinician-researchers (52%) or clinicians (45%) [n=65], and 93% reported direct involvement in deciding whether patients are offered surgery [n=68]. Just under half (46%) were surgical program directors [n=68]. Nine respondents reported they also collect, analyze and interpret the fMRI data.

Almost two thirds of respondents were from the USA (50); with further responses from Australia (6); Canada (3); France (3); Italy (3); Turkey (3); England (2); Germany (2); Denmark (1); Egypt (1); Georgia (1); Japan (1); Netherlands (1); Norway (1); Portugal (1); Scotland (1); Sweden (1); and Switzerland (1).

## 3.1 Results

### 3.2 Methods used clinically for language lateralization

Nearly all respondents reported using fMRI (96% of programs), neuropsychological assessment (99%) and extraoperative stimulation mapping (93%) in evaluating language preoperatively [total respondents: n=80] (Figure 1). The majority of programs also use intraoperative stimulation mapping (ISM) (83%), with other methods each used by fewer than half of programs. Patients are most likely to receive neuropsychological assessment (93% of patients), fMRI (58%) or Wada testing (43%) [n=80] at programs using these methods. The three programs not using fMRI (two US, one worldwide) reported they did not have fMRI capabilities, or felt it was not necessary given (e.g.) availability of Wada. Clinicians reported high confidence that, with all methods at their disposal, they can identify a patient’s language dominant hemisphere (mean 92%, SD 8; 60-100) and localize language regions to guide surgical margins (mean 84%, SD 14, 30-100) [n=81].

**Figure 1:**
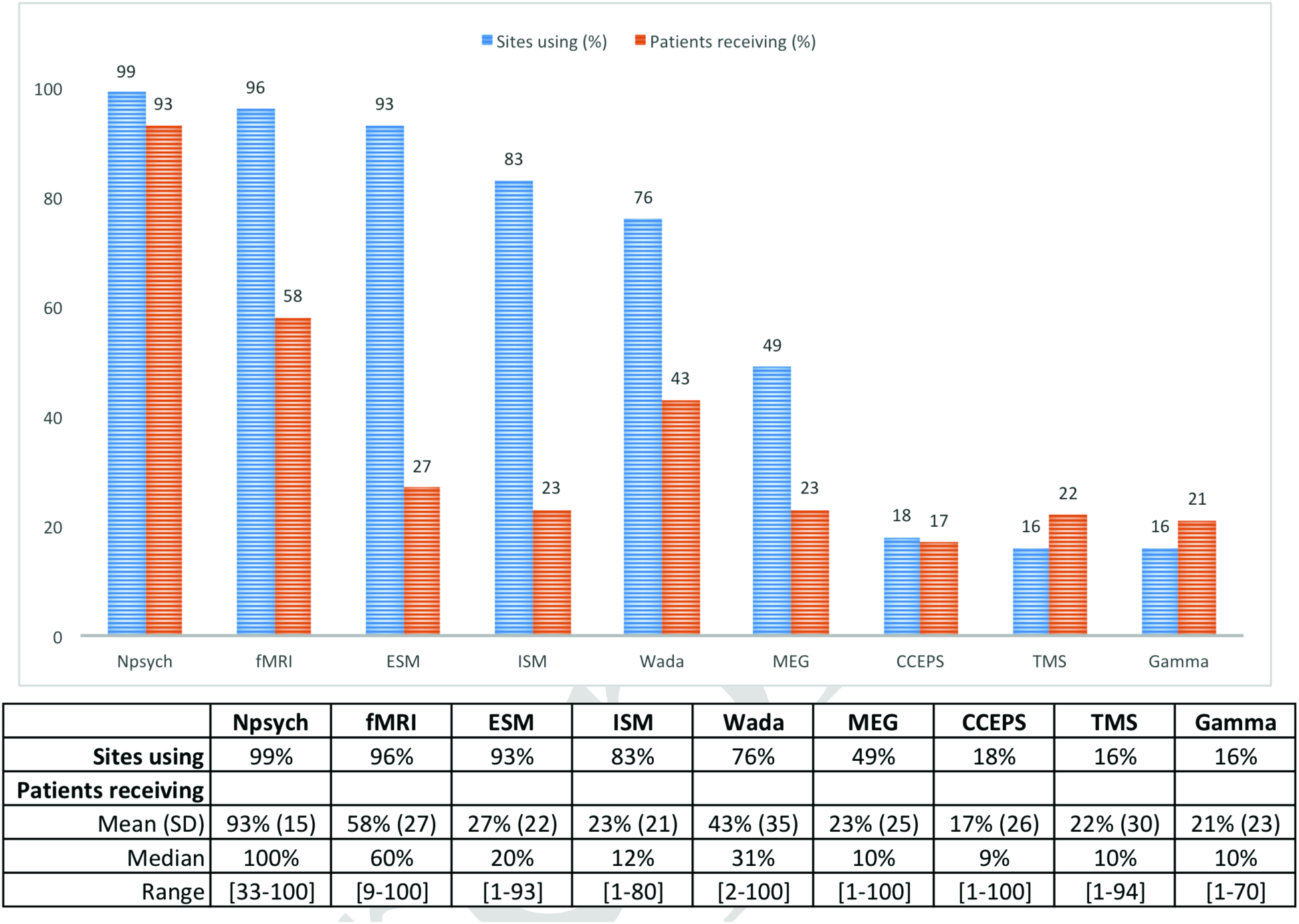
Language mapping methods: Proportion of programs using (blue) and patients receiving (orange) each method. The proportion of patients receiving a measure is estimated based only on programs using that method-i.e., at the 83% of sites using ISM, on average 23% of patients receive ISM. Neuropsychological assessment (Npsych), functional MRI (fMRI), extraoperative stimulation mapping (ESM), intraoperative stimulation mapping (ISM), Wada testing (Wada), Magnetoencephalography (MEG), Cortico-cortical evoked potentials (CCEPS), transcranial magnetic stimulation (TMS), Gamma-activation mapping (Gamma). Question: “Please estimate the proportion of surgical candidates at your center who receive the following prior to surgery to clarify language organization.”

### 3.3 Clinicians’ opinions of language fMRI

Programs using fMRI reported confidence (73%, SD 18, range 1-100) in their program’s ability to identify a patient’s language dominant hemisphere using fMRI alone [n=78]. They reported low confidence (45%, SD 27, 0-92) in their ability to use fMRI to identify specific language areas to guide surgical boundaries (i.e., localization) [n=77]. Accordingly, when considering the technique of fMRI generally they considered it reliable for language lateralization (81% confidence, SD 15 19-100) but less so for localization (i.e., identifying language areas to guide surgical boundaries; 48%, SD 25 0-93; t_(74)_=11.471 p<0.001) [n=76, 78]. When asked if language fMRI completed at different centers yields equivalent results, their response was neutral (score of 0 SD 2.5; confidence scale: strongly disagree to strongly agree [-5 to +5]).

### 3.4 Language lateralization with fMRI

All programs administering fMRI use it to identify the language dominant hemisphere [n=71]. Just under half of programs (46%) reported fMRI has never disagreed with other measures of laterality. A third of programs (34%) reported at least one instance of disagreement with Wada; a quarter (24%) with stimulation mapping; and 15% with other methods (e.g., MEG, semiology, TMS, neuropsychological testing) [n=74]. Multiple programs noted fMRI yields bilateral or equivocal findings more often than other methods, particularly in children.

In cases of disagreement fMRI was most often judged to have been incorrect (55%); a third reported cases where true lateralization remained unknown (29%); and a fifth (18%) reported cases where fMRI had been correct and the other modality was not [n=38]. Five programs reported cases of Wada-fMRI disagreement where fMRI was judged correct. None had been published, though one was included “in general in wider publications.” Of all programs reporting disagreement between fMRI and another measure, two (5%) had published these cases [n=39].

### 3.5 Language localization with fMRI

Programs reported using fMRI to localize language less frequently (44%) [n=71].

Most centers using fMRI to localize language (guide surgical margins and preserve language cortex) seek to map known language areas (86%) [n=29]. These include Broca’s (86%) and Wernicke’s areas (76%), with "basal temporal language area" (BTLA; 17%) and other regions (21%) mapped infrequently (e.g., Supplementary Motor Area; SMA).

Programs that do not use fMRI to localize language (i.e., programs using fMRI only for language lateralization) less often seek to map known language regions (58%) (Fisher’s exact p<0.001) [n=40]. Specifically, these programs map Broca’s (43%; p<0.001) and Wernicke’s Areas (38%; p=0.003) less frequently than programs using fMRI for localization, and also rarely map BTLA (5%, p=0.122) or other regions (5%; p=0.062).

Clinicians using fMRI to localize language report success in mapping Broca’s 75% of the time (SD 15%, 39-93%) [n=25]; in mapping Wernicke’s 71% of the time (SD 16%, 30-92%) [n=22]; BTLA 62% of the time (SD 28%, 19-90%) [n=5] and other regions 52% of the time (SD 31%, 3-80%) [n=6]. The few who use fMRI for localization, but do not seek to identify specific language regions [n=4], find fMRI is rarely successful (27% of cases) in guiding margins.

Direct electrical stimulation at fMRI-positive cortex typically confirms fMRI findings (74% of the time; SD 15%, 30-95%), with most of these respondents (86%) reporting they specifically seek to map certain language regions [n=28]. Instances of post-operative language decline were reported with equivalent frequency (26%) by both programs using fMRI for lateralization only [n=27] and those using it for both lateralization and localization [n=23].

### 3.6 Cognitive outcomes when fMRI activation is preserved

When no fMRI-positive language cortex is resected, half of responding programs (49%) reported no cases of persisting language decline three months post-surgery [n=75]. Seventeen percent reported at least one otherwise unexplained instance of language decline following temporal [67%], frontal (50%) or less often parietal (33%) or occipital (8%) surgery [n=12]. Of note, most patients (70%) receive postoperative neuropsychological testing at most (96%) programs [n=79]. None of these cases of decline had been published. A third of programs did not know if cases of decline had occurred.

Programs noting cases of decline most often related these to surgical variables (100% reported at least one such case) and 23% reported instances where fMRI-related factors were considered responsible. Surgically, resection of white matter language tracts was most often noted (85%). Resection too close to fMRI-positive areas was also often reported (38%), as were surgical complications (23%); multiple subpial transections over eloquent cortex (15%); and surgical injury outside the planned resection area (15%). Resection in language association cortex and post-surgical inflammation was also noted, and 31% reported cases where the cause was unknown. With respect to fMRI-related factors, different programs attributed at least one instance of unanticipated deficits to imaging (e.g., artifact, insufficient resolution) (n=1); analysis (n=1); and patient-related factors (e.g., seizures, movement) (n=1).

In exploratory analysis, programs reporting cases of cognitive decline in spite of fMRI-active areas being preserved did not differ from those which did not on a range of variables (Supplement 3). While a nonsignificant difference, in this sample programs reporting unexpected decline more often predominantly evaluated children (31% v 11%); were less likely to review the actual fMRI images at conference (69% v 89%); and less often had the individual involved in fMRI data analysis interpret data at surgical conference (38% v 59%).

### 3.7 Cognitive outcomes when fMRI activation is resected

Half of the programs using fMRI to guide surgical margins (54%) reported cases where patients maintained function after resection of language-positive sites [n=26]. These followed frontal (62%), temporal (54%), or parietal (23%) resections. Select sites noted surgery in the left operculum (bilaterally active), posterotemporal cortex (unspecified); right posterotemporal cortex (bilateral activation) and middle frontal gyrus. None had been published.

In exploratory analysis, these programs were less likely to use the Wada (in at least some instances) to evaluate language organization (Supplement 3), while use of language fMRI was equivalent (61% (SD 25) vs 64% (SD 24) [n=14,12]). Other nonsignificant differences in this sample included programs reporting unexpected preservation more often loading images into an intraoperative system (64% vs 33%); using fMRI to localize language and guide surgical margins (57% vs 33%); and completing more surgeries annually (42 (SD 21) vs 34 (SD 16)).

### 3.8 How clinical teams interpret fMRI maps

Teams who routinely localize language areas with fMRI to guide surgical margins most often constrain resections to a given distance from fMRI activation (59% of programs) [n=29]. This distance threshold used by programs varied from 3-50mm (average 15mm, SD 12). The most commonly-used margin was 10mm (42% of programs). Programs often (28%) will not operate in the same gyrus as activation or use other criteria (28%), most often routine investigation of activation with direct stimulation. These programs most often report they would resect fMRI-positive cortex in at least some circumstances (79%), for instance if cleared by direct stimulation (62%); if activation was not anatomically consistent with a language area (38%); if deficits would likely be temporary (e.g., uncomplicated unilateral SMA resection) (38%); or if the patient was willing to accept post-surgical deficits (24%) [n=29]. One site noted they would resect any fMRI-positive activation “not [in] a primary language site (Broca/Wernicke)” (this site reported both unpredicted decline and unexpected preservation), while one site was uncertain, and 17% reported they would never resect cortex that was fMRI language-positive.

To interpret fMRI, teams most often review the images (visually) at surgical conference (86% of programs) and the team or referrer reviews a written report (71%) (n=79). The individual involved in analysis often interprets the data for the team (63%). Laterality indices are rarely used (whole brain, 28%; specific region, 18%) and a third of programs load data into an intraoperative mapping system (35%).

Respondents expressed confidence that fMRI’s ability to lateralize language will improve with further technical advances (mean=89%, SD 12) though were less confident about future advances improving fMRI’s utility for localization (mean=64%, SD 26%; n=77) (t_(75)_=8.831, p<0.001).

## 4.1 Discussion

These results confirm fMRI is well established for language lateralization, and is already used in many epilepsy programs to localize language cortex to help guide surgical margins. Consistent with the research literature,^6^ most clinical programs using language fMRI (54%) reported instances where fMRI language lateralization had conflicted with other methods including the Wada (34%). In these situations, it is rarely clear which method is “correct,” though when they could do so respondents judged fMRI incorrect more often (55%). These findings may suggest the fMRI tasks used by different programs vary in their ability to predict language outcome; reflect a decline in language not captured by visual naming tasks;^15^ or represent recall bias and clinicians’ uncertainty regarding fMRI.

While fMRI is not validated for language localization, programs have already begun to cautiously use it for this purpose (44%), typically removing fMRI-positive cortex only after confirmation via direct stimulation, counseling patients on possible decline, or never removing fMRI-positive cortex. This emphasizes the importance of using and further validating highly standardized forms of fMRI.^12^ Cases of unexpected preservation and unexpected decline are equally concerning; the former may increase post-surgical cognitive impairment while the latter may increase surgical failure rates.^16^ Until validated, any criteria used to "preserve" functional cortex relative to fMRI map boundaries in surgery are arbitrary.

When interpreting fMRI maps, it should be emphasized that rather than showing the immutable boundaries of language-critical cortex, the map reveals the probable location of language areas. The language map also represents assumptions made by the individuals who selected the cognitive task and imaging sequences, and analyzed and reported the data. These assumptions can dramatically change map boundaries. Examples are shown in Figure 2. The perceived boundaries of language areas will vary with a patient’s language skill (Figure 2A); with analysis parameters (3B); with signal loss that is not evident in the final map (3C); with the task and control conditions used (3D); and the areas the analyst is asked (or not asked) to identify (3E). The latter is of particular significance given clinicians’ current focus on Broca’s and Wernicke’s but not other known language-critical regions (e.g., BTLA, Exner’s area, SMA).^17-21^ Just as importantly, language fMRI represents task-correlated changes in blood oxygenation (rather than neural activation) and may show task-correlated but not critical regions.^22^

**Figure 2:**
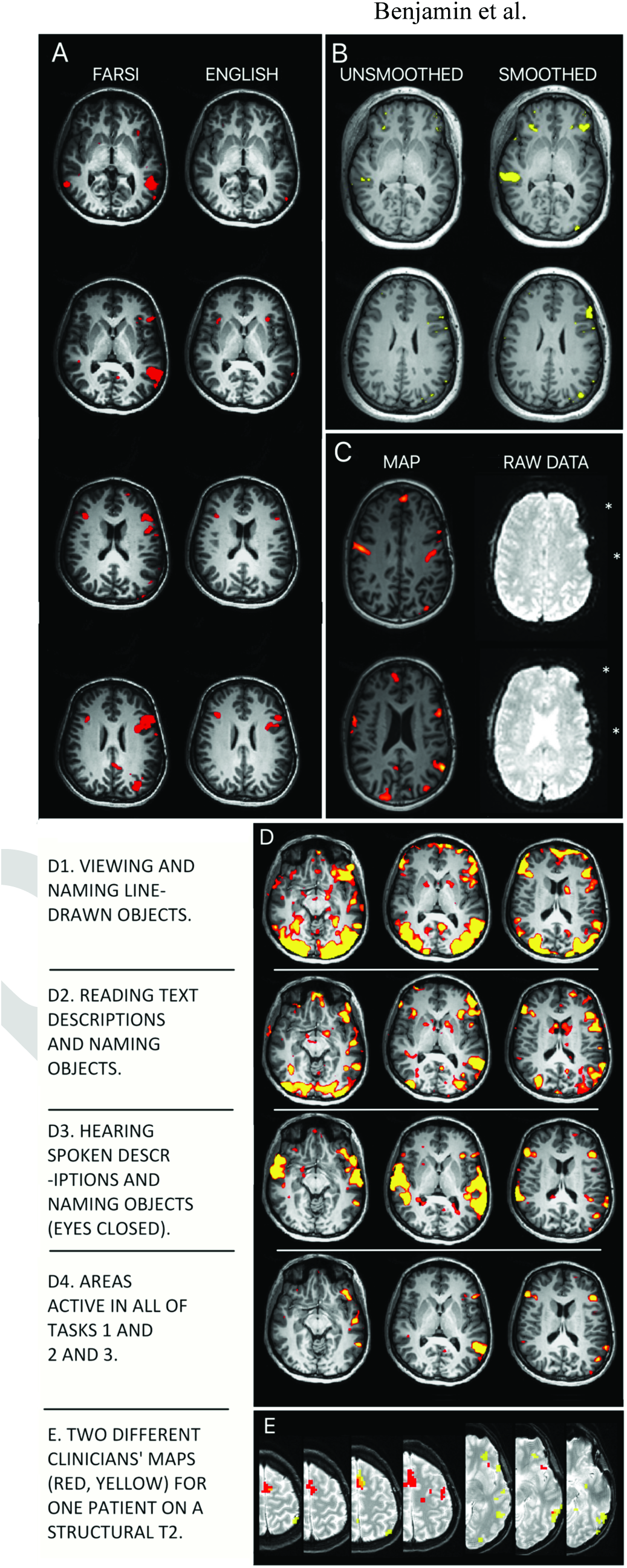
Language maps will vary in clinically meaningful ways due to multiple variables. Surgical teams can manage these factors by using experts in both imaging (e.g., radiology) and cognition (e.g., neuropsychology) in clinical fMRI design, analysis and interpretation. <u>A: Language skill</u>. A patient’s language ability will change their activation map. Maps using the same tasks in Farsi and English from a patient who reported fluency in, and made medical decisions in English. <u>B: Data analysis</u>. Each analysis step changes the map. Data "smoothing" removes noise. Whether it is appropriate, and to what degree, is debated. The degree of smoothing in commercial software may be unspecified. Identical analysis without (left) and with (right) smoothing (8mm kernel). <u>C: Data quality</u>. A statistical map (left) does not show where raw data are missing (right, asterisks). These areas will not be active even if they are language critical. This map (left) was presented to a surgical team for surgical planning without caveat. <u>D. Cognitive task</u>. Different language tasks give different maps. Subtle changes in task instructions, patient motivation and cognitive strategy change language maps. (D1) Visual object, (D2) text reading and (D3) auditory tasks are shown as well as (D4) the intersection of these. <u>E: Analyst expectations</u>. The analyst’s perceived goal will change the activation map. Two overlaid maps (red; yellow) generated independently by two clinicians for the same patient (see Benjamin et al., 2017).^21^ Analysts were blind to case details. One prioritized frontal (red) and the other temporal (yellow) regions, as when mapping frontal tumor vs temporal lobectomy cases. Overlap in orange. 3E is reprinted from Benjamin et al., "Presurgical language fMRI: mapping of six critical regions," Human Brain Mapping (2017); Creative Commons Attribution-Non-Commercial License.

When cognitive tasks and analysis packages are received from MR vendors and used without detailed consideration, how many key variables are addressed is opaque. US clinicians are not mandated to use FDA approved software, and delegating these considerations is risky when perhaps the best validated language protocols and analysis software to date are open-source and freely available (Supplement 2). The clinician is ultimately responsible for their results. The skills required for clinical fMRI (selecting sequences and cognitive paradigms; analyzing and interpreting data) do not fit within one existing discipline. When a gold standard develops, it will likely approximate the interdisciplinary Wada protocol with a team of at least two qualified professionals from a selection of radiology, neuropsychology, neuroscience, and/or engineering. Expertise in the clinical, imaging and cognitive skills required for fMRI, rather than an individual’s discipline, will likely best predict quality of care.

The above discussion fits with the observation that, in our sample, programs which reported unexpected decline were less likely to review the data in surgical conference (69% v 89%) and less likely to have the individual who completed analysis interpret data at conference (38% v 59%; though note that neither reached statistical significance). The greater use of the Wada for language lateralization by programs not reporting unexpected decline, despite their using language fMRI as regularly as other programs, may reflect a more cautious supplementation of fMRI with Wada when results are equivocal or uncertain.

## 4.2 Limitations

A critical limitation of this report is our inability to relate these findings to the specific language tasks used. Of note, variation in protocols is likely to influence overall network lateralization less than regional localization. It is possible respondents’ recall and reports are imperfect, and it is not possible for us to judge what reports of unexpected decline or preservation are based on (e.g., formal testing, patient report). These data better represent American (61%) and academic (85%) epilepsy programs. To increase our response rate, collaborators forwarded the survey to numerous colleagues, obscuring the true response rate. As an estimate, we contacted 221 US NAEC programs and received 50 US responses, suggesting a 23% response rate (using this approach Hamberger et al.^13^ received 39 US responses). The survey length will also have decreased responses.

## 4.3 Conclusions

Clinical fMRI is widely used to predict language laterality and post-surgical language change. The caveats documented in the literature-e.g., occasional disagreement with Wada-are seen in the clinic. Outcomes can likely be improved through use of existing, well validated tasks. Our data, from centers using a range of tasks and methods, emphasize that cautious use of language fMRI for lateralization is warranted and that fMRI maps cannot simply be treated as representing language-critical cortex. Without standardization and explicit validation, any criteria for preserving language-critical cortex relative to fMRI map boundaries are arbitrary. We suggest an initial minimum for clinical care might involve ensuring those who analyze a program’s language fMRI, and understand the task’s cognitive design, interpret the data in 3D in discussion with the team at conference. They might begin by reviewing the task itself (expected activation), the best estimates of its sensitivity and specificity, and the limitations of the specific results (patient factors, areas of signal loss). This will reduce opportunity for misinterpretation and likely improve patient care.

## Acknowledgement

Thank you to our respondents: this required significant time and was a generous contribution.

## Supplement 1

The survey questions completed by respondents.

Note that text in square brackets is added here for clarity. Please note that whenever respondents used a sliding bar to respond (“slider”), the precise value they were selecting was displayed in increments of 1 point. As such, when a slider had given numbers above it for reference (e.g., a scale of 0, 10, 20… 100) the respondent’s selection still reflected a precise response (e.g., 61, 93, etc.). Notes regarding not attending AES 2015, where preliminary data were presented, were added at that time before the final few international respondents added their responses.

<u>The Yale Survey of Clinical Language fMRI in Epilepsy</u>

Should I complete this questionnaire?

We are asking for responses from both:

1. The director of the epilepsy surgical program, or a senior clinician involved in determining patients’ surgical eligibility; and:
2. The individual involved in collecting, analyzing, and potentially billing clinical language fMRI for surgical planning at your institution.

If you are one of the above, please continue. There will be a link for the other relevant person at the end of your survey. If you are not one of the above, we would be grateful if you could forward the link to the relevant person at your organization.

Q1 **Informed Consent** Thank you for your help! While your responses will not be anonymous, they will not be reported in a way that specifically links you to your responses. Data will only be reported in an aggregate or de-identified manner. Only the researchers involved in this study and those responsible for research oversight will have access to the information you provide. We do not ask for and request you do not include any patient information or other sensitive information. Participation in this study is completely voluntary. You are free to decline to participate, to end participation at any time for any reason, or to refuse to answer any individual question at any time. There are no direct benefits or known risks to you for participating. If you have any further questions about this study, you may contact Alexa Li at or Dr. Christopher Benjamin at . If you would like to speak with someone other than the researchers to discuss problems, concerns, and questions you may have concerning this research you may contact the Yale Human Investigation Committee at +1 (203) 785-4688.

- I consent. I did not attend the AES Neuropsychology SIG and have not seen this study’s preliminary data. (1)

Q2 We greatly appreciate your time; you are one of the few people who can help us run this study. Our priority is balancing your time with obtaining critical information to improve clinical language fMRI. In piloting, this survey took respondents 7-10 mins. There is one set of questions if you are a Surgical Director (decides whether patients are offered surgery); another if you collect or analyze language fMRI data. Please note: 1. If you would like to receive our preliminary, pre-publication findings (a video presentation) please indicate on the final survey page (available late 2015). 2. Estimate responses (or use “don’t know” if available) if obtaining exact answers is difficult. 3. Nearly all questions are optional. Non-optional questions have a “Don’t know” choice.

Q3 To your knowledge, has someone in your program already completed part of this survey and forwarded you a link?

- Yes (1)
- No (2)

*[Answered If: Q3 To your knowledge, has someone in your program already completed part of this survey and forwarde…Yes Is Selected]*

Q4 Please enter the survey code your colleague sent you (6-digit number). We cannot use your data without this code to link your responses.In the following, please select the set of questions (“Director” or “Technical”) noted in the email you received.

Q5 What is your role in your epilepsy surgical program? This determines what questions you answer. Please select one. Most individuals do not do both.

- A. I select patients for neurosurgery. I do not personally collect or process fMRI data (e.g., epilepsy surgical program director, neurosurgeon, epileptologist) (1)
- B. I personally collect, analyze, interpret clinical fMRI data. I use software like SPM (e.g., radiologist, neuropsychologist, imaging scientist etc.) (4)
- C. I do both. Select with caution (see note above). (3)

[Respondents who selected 5B completed a different version of this survey and are not included here].

Q6 The following relate to decision making in your surgical program, with the standard clinical language fMRI available to you for this purpose. To what extent do you agree with the statements-

Q7 “Our program can identify a patient’s language dominant hemisphere…”

______ “…using all methods at our disposal” (not just fMRI) (1)

______ “…using fMRI alone” (2)

*[Respondents used a slider to respond, marked from 0 (not at all confident) to 100 (highly confident) numbered in increments of 10 - i.e., 0, 10 20 30…]*

Q8 “Our program can identify specific language regions to guide surgical margins…”

______ “…using all methods at our disposal” (not just fMRI) (1)

______ “…using fMRI alone” (2)

*[Respondents used a slider to respond, marked from 0 (not at all confident) to 100 (highly confident) numbered in increments of 10 - i.e., 0, 10 20 30…]*

Q9 The following questions relate to clinical language fMRI as a general technique, completed at any center worldwide.

Q10 “The technique of fMRI, is a reliable technique for identifying…”

______ “…the language dominant hemisphere.” (1)

______ “…language areas to guide surgical boundaries.” (E.g. Broca’s, Wernicke’s) (2)

*[Respondents used a slider to respond, marked from 0 (not at all confident) to 100 (highly confident) numbered in increments of 10 - i.e., 0, 10 20 30…]*

Q11 “With further technical advances, the technique of fMRI will eventually reliably identify patients’…

______…language dominant hemisphere” (1)

______…language areas to guide surgical boundaries.” (E.g. Broca’s, Wernicke’s) (2)

*[Respondents used a slider to respond, marked from 0 (not at all confident) to 100 (highly confident) numbered in increments of 10 - i.e., 0, 10 20 30…]*

Q12 To what extent do you agree that:

Q13 “Language fMRI as currently completed at different centers yields equivalent results.”

______(1)

*[Respondents used a slider to respond, marked from −5 (strongly disagree) to 0 (Neutral) to 5 (strongly agree) numbered in increments of 1 - i.e., −5 −4 −3…]*

Q14 “Language fMRI is useful in predicting post-surgical language outcome.” (impairment or residual functional capacity)

______(1)

*[Respondents used a slider to respond, marked from −5 (strongly disagree) to 0 (Neutral) to 5 (strongly agree) numbered in increments of 1 - i.e., −5 −4 −3…]*

Q60 Is your epilepsy program affiliated with a university?

- Yes (1)
- No (2)

Q61 Please estimate the number of patients, per year, who your program:

______evaluates for surgical candidacy (1)

______treats surgically (2)

*[Respondents used a slider to respond to each of these, marked from 0 to 300 in increments of 50]*

Q62 Your program predominantly evaluates:

- Children ((1)
- Adults (2)
- Both (3)

*[Respondents could only select one of these]*

Q63 Please estimate the proportion of surgical candidates at your center who receive the following prior to surgery to clarify language organization?

______Language Functional MRI (1)

______Wada testing (2)

______Extraoperative cortical stimulation mapping (grids, strips) (3)

______Intraoperative cortical stimulation mapping (4)

______MEG (5)

______TMS (6)

______Neuropsychological assessment (8)

______Gamma activation mapping (10)

______CCEPS (11)

______Other (please specify): (7)

______Other 2 (please specify): (12)

*[Respondents used a slider labelled “Percentage (%)” to respond, marked from 0 to 100 numbered in increments of 10 - i.e., 0 10 20 30…]*

Q71 Which of the following language fMRI data does your team use? (Select all that apply)

- Team or referrer reviews written report (1)
- Team reviews images visually at surgical conference (2)
- A laterality index is used, based on select regions of interest (3)
- A laterality index is used, based on whole brain activation (4)
- Images loaded into intraoperative mapping system (e.g. Stealth) (5)
- The individual involved in analysis interprets the data at surgical conference. (7)
- Other (6)____________________

*[Respondents could check multiple of these]*

Q72 Of those patients who undergo epilepsy surgery, what portion receive post-operative neuropsychological follow up (estimated)?

______(1)

*[Respondents used a slider to respond, marked from 0 to 100 numbered in increments of 10 – i.e., 0 10 20 30…]*

Q73 Does your institution routinely use fMRI to (click all that apply):

- Determine the language dominant hemisphere (1)
- Guide surgical margins to avoid language cortex (2)

*[Respondents could check multiple of these]*

Q74 In your program’s actual experience, approximately how often do you consider language fMRI to have successfully identify the language dominant hemisphere?

______% Success (1)

*[Respondents used a slider labelled “% Success” to respond, marked from 0 to 100 numbered in increments of 10 - i.e., 0 10 20 30…]*

Q75 Has fMRI-defined language laterality ever disagreed with laterality determined by other measures?

- Yes, disagreed with Wada result (1)
- Yes, disagreed with direct stimulation mapping (2)
- Yes (other) (3)____________________
- No (4)

*[Respondents could check multiple “Yes” options, or No]*

*[Answered If: Q75 Has fMRI-defined language laterality ever disagreed with laterality determined by other measures? No Is Not Selected]*

Q76 When fMRI-defined language laterality disagreed with results from a measure we/I concluded: (Can select multiple if multiple cases)

- fMRI lateralization was correct. (1)____________________
- fMRI lateralization was incorrect (2)____________________
- unknown. (3)____________________

*[Respondents could check multiple options]*

*[Answered If: Q75 Has fMRI-defined language laterality ever disagreed with laterality determined by other measures? No Is Not Selected]*

Q77 Have you published this case(s)?

- Yes (optional: please provide author, year or citation) (1) ____________________
- No (3)____________________

Q78 Using fMRI, do you routinely systematically map the boundaries of:

- Broca’s (1)
- Wernicke’s (2)
- Basal temporal naming areas (3)
- Other language areas (4)__________________
- No (5)

*[Respondent can select multiple of 1-4, or can select 5 alone]*

*[Answered If: Q78 Using fMRI, do you routinely systematically map the boundaries of: No Is Not Selected]*

Q79 In your program’s clinical experience, approximately how often does language fMRI successfully help you guide surgical margins by mapping the boundaries of:

______Broca’s area (1)

______Wernicke’s area (2)

______Basal Temporal Naming Areas (3)

______Other areas (e.g., SMA) (5)

*[Respondents used a slider labeled “Percentage (%)” to respond, numbered from 0 to 100 in increments of 10 - i.e., 0 10 20 30…]*

*[Answered If: Q78 Using fMRI, do you routinely systematically map the boundaries of: No Is Selected]*

Q80 In your program’s clinical experience, approximately how often does language fMRI successfully guide surgical margins of language?

______Percentage (%) (1)

*[Respondents used a slider labeled “Percentage (%)” to respond, numbered from 0 to 100 in increments of 10 - i.e., 0 10 20 30…]*

*[Answered If: Q73 Does your institution routinely use fMRI to (click all that apply): Guide surgical margins to avoid language cortex Is Selected]*

Q81 What percentage of cases who complete language fMRI to guide resection margins also receive cortical stimulation mapping (intra-or extraoperative)?

______Percentage of cases (none=0) (1)

*[Respondents used a slider numbered from 0 to 100 in increments of 10 - i.e., 0 10 20 30…]*

*[Answered If: Q81 What percentage of cases who complete language fMRI to guide resection margins also receive corti…Percentage of cases (none=0) Is Greater Than 0]*

Q82 In these cases, how often do you estimate fMRI-positive language areas are also positive on direct stimulation?

______Percentage (%) (1)

*[Respondents used a slider numbered from 0 to 100 in increments of 10 - i.e., 0 10 20 30…]*

*[Answered If: Q73 Does your institution routinely use fMRI to (click all that apply): Guide surgical margins to avoid language cortex Is Selected]*

Q83 Please complete: To avoid post-surgical language deficits, we will not operate within -(note: select all that apply)

- A given distance of a language activation on fMRI. (1)
- The same gyrus as a language activation on fMRI. (2)
- We would resect an fMRI positive language area. (5)
- Other (4)________________

*[Respondents could select multiple]*

*[Answered If: Q83 Please complete: To avoid post-surgical language deficits, we will not operate within - (note: se…A given distance of a language activation on fMRI. Is Selected]*

*[Note also these respondents must also have indicated that they use fMRI to guide surgical margins to avoid language cortex in Q73]*

Q84 When using language fMRI to guide resection margins, what is the closest distance to an activation that you will operate?

______millimeters (1)

*[Respondents used a slider labeled “millimeters” and numbered from 0 to 50 in increments of 5 - i.e., 0 5 10…]*

*[Answered If: Q73 Does your institution routinely use fMRI to (click all that apply): Guide surgical margins to avoid language cortex Is Selected]*

Q85 In surgery, does your team ever remove fMRI-identified, language positive sites? (Select all that apply)

- Never (1)
- Yes, if the area is cleared using direct stimulation (2)
- Yes, if we do not believe anatomically it is a language area (3)
- Yes, if any deficit will most likely be temporary (e.g., uncomplicated unilateral SMA) (4)
- Yes, if the patient is willing to accept a post-surgical language deficit (7)
- Yes, other (5)____________________
- Don’t know (6)

*[Respondents could select multiple “Yes” and “Don’t know” options; or alternately could select only “Never”]*

*[Note: All respondents received the following question]*

Q86 So far as you know, have any patients experienced a persistent (>3 month) postoperative language decline in spite of your preserving all fMRI-positive language sites?

- Yes (1)
- No (2)
- Don’t know (3)

*[Only one response could be selected]*

*[Answered if: Q86 So far as you know, have any patients experienced a persistent (3 month) postoperative language…Yes Is Selected]*

Q87 Why do you believe this occurred? (select all that apply)

- Resection too close to fMRI-identified language positive site (1)
- Resection included white matter language tracts (2)
- Multiple subpial transections over eloquent cortex (3)
- Unexpected surgical complication (4)
- Surgical injury outside of the planned resection area (10)
- fMRI mapping was inadequate due to: (5)

*Unable to completely map resection area using fMRI due to:*

- Patient factors (e.g., seizures or movement during imaging) (6)
- fMRI image factors (e.g,. image artifact, insufficient resoultion) (7)
- fMRI analysis factors (e.g., analysis suboptimal) (8)
- I do not know (16)
- Other (please specify): (9)____________________

*[Can select multiple]*

*[Answered If: Q86 So far as you know, have any patients experienced a persistent (3 month) postoperative language…Yes Is Selected]*

Q88 If known, what area(s) were resected in these case(s)? (optional: gyrus, location)

- Frontal lobe (1)____________________
- Temporal lobe (2)____________________
- Parietal lobe (3)____________________
- Occipital lobe (4)_____________
- Don’t know (5)

*[Can select multiple]*

*[Answer If: Q86 So far as you know, have any patients experienced a persistent (3 month) postoperative language decline in spite of your preserving all fMRI-positive language sites? Yes Is Selected]*

Q89 Have you published this case/any of these cases?

- Yes (optional: please provide author, year or citation) (1)____________________
- No (3)____________________

*[Answer If: Q85 In surgery, does your team ever remove fMRI-identified, language positive sites? (Select all that…) Never Is Not Selected]*

Q90 As far as you know, have any patients maintained pre-operative language ability despite resected fMRI-positive language cortex?

- Yes (1)
- No (2)
- Don’t know (3)

*[Answer If: Q90 As far as you know, have any patients maintained pre-operative language ability despite resected fMRI-positive language cortex Yes Is Selected]*

Q91 If known, where was the fMRI-positive area(s) resected without post-operative deficit (e.g., “posterior inferior temporal gyrus”).

- Frontal lobe (1)____________________
- Temporal lobe (2)____________________
- Parietal lobe (3)____________________
- Occipital lobe (4)____________________
- Don’t know (5)

*[Can select multiple]*

*[Answer If: Q90 As far as you know, have any patients maintained pre-operative language ability despite resected fMRI-positive language cortex Yes Is Selected]*

Q92 Have you published this case?

- Yes (optional: please provide author, year or citation) (1)____________________
- No (3)____________________

*[Respondents who indicated in question 5 that they also analyze language fMRI data were then presented with a set of other questions on fMRI data acquisition and analysis before continuing.]*

Q93 Before completing the final two pages of this survey,

**If the person who acquires and analyzes fMRI data has not completed their survey please forward them the below link and key now. We cannot use your data without their responses as well. Please copy and paste the following. Thank you!**________________

When you have a chance could you please complete the 7-10 minute Yale Surgical fMRI survey for: B. The professional who collects and analyzes fMRI data? Thank you! It is at:

1. Link: http://bit.ly/yale_invitation
2. Survey key: *[A unique identifier was generated, and documented]*

Q94 I am a -

- MR technologi st (1)
- Neurologist (2)
- Neuropsychologist (3)
- Neuroscientist (9)
- Neurosurgeon (5)
- Physicist/Engineer (6)
- Radiologist (7)
- Other (8)

*[One option could be selected.]*

Q95 I currently work as a -

- Clinician (1)
- Researcher (2)
- Both (3)

*[One option could be selected.]*

Q96 Are you directly involved in deciding for your surgical program whether patients are offered surgery?

- Yes, I am the Surgical Program Director (2)
- Yes, though I’m not the Surgical Program Director (4)____________________
- No, in our program I am the - (3)____________________

*[One option could be selected. Option 3 allowed for a free text response.]*

Q106 (Optional): Our hospital/Institution is:

*[Text response could be entered]*

Q97 Please provide your contact information if:

1. You would like us to email you a link to a youtube video with our findings prior to publication.
2. We can contact you to clarify questions
3. You would like access to a communal site hosting details on clinical fMRI paradigms, sequences, etc.

Name: (1)

Email: (2)

*[Text response could be entered]*

Q98 Comments:

*[Text response could be entered]*

Q99 Go to the final page to submit and you’re done!

## Supplement 2

The semantic decision-making language task discussed in text is available directly from Jeff Binder. A version of the task, using the same stimuli within “Presentation” software, is also freely available at cogneuro.net/epilepsia2017. The analysis software noted includes (is not limited to) SPM (www.fil.ion.ucl.ac.uk/spm/) and FSL (fsl.fmrib.ox.ac.uk/fsl) are excellent, open, and well documented software packages for fMRI analysis that are widely used that are continually refined and improved. <u>Note on FDA approval and fMRI software</u>: In the US, for clinical care, the clinician completing analysis is responsible for determining which software they use, and how they use it. Their choice is not constrained to US Food and Drug Agency (FDA) approved products; the FDA constrains what the marketers of software state the software should be used for. That is, if software is FDA approved for a given purpose, the FDA has judged it can be used for this purpose.

## Supplement 3

<u>3A. Characteristics of programs reporting unexpected language decline and unexpected language preservation. Exploratory analysis</u>. Note that sample size varies as a function of the number of respondents answering all required questions. Unexpected decline: All respondents were asked “So far as you know, have any patients experienced a persistent (>3 month) postoperative language decline in spite of your preserving all fMRI-positive language sites?” (Q86). Unexpected preservation: The question “as far as you know, have any patients maintained preoperative language ability despite resected fMRI-positive language cortex?” (Q90) was shown to all respondents who did not report they would never remove fMRI-identified, language positive sites (Q85). DCS = Direct Cortical Stimulation mapping.

| Unexpected decline. |  |  |  |  |  |
| --- | --- | --- | --- | --- | --- |
| Program characteristics | Decline | No Decline | p<sup>^</sup> | N (respondents) |  |
|  |  |  |  | Yes | No |
| Average number of patients evaluated annually | 116 (83) | 113 (70) | 0.897 | 13 | 37 |
| Number of surgical cases annually | 38 (24) | 34 (21) | 0.524§ | 13 | 37 |
| Proportion of patients who also receive DCS<sub>j</sub> | 59% | 59% | 0.977 | 6 | 17 |
| Program never uses Wada to clarify language org'n | 23% | 32% | 0.728 | 13 | 37 |
| Program is willing to resect fMRI-positive areas* | 60% | 88% | 0.210 | 5 | 17 |
| Median resection margin cf. fMRI (mm)• | 10mm | 10mm | — | 5 | 7 |
| Program predominantly evaluates children, adults° | 31%, 23% | 11%, 46% | 0.142 | 13 | 37 |
| Program is university-affiliated | 92% | 84% | 0.660 | 13 | 37 |
| Interpretation and use of fMRI data |  |  |  |  |  |
| Programs using fMRI to localize language areas | 55% | 49% | 1.000 | 11 | 35 |
| Team reviews a written report | 69% | 65% | 1.000 | 13 | 37 |
| Team reviews images at conference | 69% | 89% | 0.181 | 13 | 37 |
| Programs using a laterality index – Select region | 23% | 32% | 0.728 | 13 | 37 |
| Programs using a laterality index – Whole Brain | 8% | 19% | 0.662 | 13 | 37 |
| Programs loading images into intraoperative system | 31% | 41% | 0.742 | 13 | 37 |
| Individual who analyzes the data presents it | 38% | 59% | 0.215 | 13 | 37 |

<sup>^</sup>Fisher's Exact or t-test as appropriate; <sub>j</sub>The proportion of patients who also receive stimulation mapping, at sites who localize language; \*e.g., if cleared by stimulation mapping; °Sites reporting they predominantly evaluate "both" children and adults excluded. •The mean of 10mm was reported by 3 sites in each of those (i) reporting and (ii) not reporting false negatives.

^Fisher’s Exact or t-test as appropriate; ¡The proportion of patients who also receive stimulation mapping, at sites who localize language; *e.g., if cleared by stimulation mapping; ºSites reporting they predominantly evaluate “both” children and adults excluded. •The mean of 10mm was reported by 3 sites in each of those (i) reporting and (ii) not reporting false negatives.

<u>3B. Characteristics of sites reporting at least one patient who had an unexpected preservation of language function in spite of fMRI-identified language areas being resected. Exploratory analysis</u>. Specifically, the question “as far as you know, have any patients maintained pre-operative language ability despite resected fMRI-positive language cortex?” (Q90) was shown to all respondents who did not state that they would never remove fMRI-identified, language positive sites (Q85).

| Unexpected preservation. |  |  |  |  |  |
| --- | --- | --- | --- | --- | --- |
| Program characteristics | Preserved | Not Preserved | p<sup>^</sup> | N (respondents) |  |
|  |  |  |  | Yes | No |
| Average number of patients evaluated annually | 114 (54) | 105 (58) | 0.671 | 14 | 12 |
| Number of surgical cases annually | 42 (21) | 34 (16) | 0.256 | 14 | 12 |
| Proportion of patients who also receive DCS<sub>j</sub> | 58% (26) | 69% (36) | 0.612 | 8 | 4 |
| Program never uses Wada to clarify language org'n | 42% | 0% | 0.020 | 14 | 11 |
| Program is willing to resect fMRI-positive areas* | 100% | 100% | – | 8 | 4 |
| Median resection margin cf. fMRI (mm)• | 5, 8, 10, 50 | 4,10 | – | 4 | 2 |
| Program predominantly evaluates children, adults° | 29%, 36% | 17%, 42% | 0.633 | 9 | 7 |
| Program is university-affiliated | 93% | 92% | 1.000 | 14 | 12 |
| **Interpretation and use of fMRI data** |  |  |  |  |  |
| Programs using fMRI to localize language areas | 57% | 33% | 0.267 | 14 | 12 |
| Team reviews a written report | 64% | 83% | 0.391 | 14 | 12 |
| Team reviews images at conference | 93% | 83% | 0.580 | 14 | 12 |
| Programs using a laterality index – Select region | 43% | 25% | 0.429 | 14 | 12 |
| Programs using a laterality index – Whole Brain | 14% | 25% | 0.635 | 14 | 12 |
| Programs loading images into intraoperative system | 64% | 33% | 0.238 | 14 | 12 |
| Individual who analyzes the data presents it | 64% | 50% | 0.692 | 14 | 12 |

<sup>^</sup>Fisher's Exact or t-test as appropriate; <sub>j</sub>The proportion of patients who also receive stimulation mapping, at sites who localize language; \*e.g., if cleared by stimulation mapping; °Sites reporting they predominantly evaluate "both" children and adults excluded. •The mean of 10mm was reported by 3 sites in each of those (i) reporting and (ii) not reporting false negatives.

^Fisher’s Exact or t-test as appropriate; ¡The proportion of patients who also receive stimulation mapping, at sites who localize language; *e.g., if cleared by stimulation mapping; ºSites reporting they predominantly evaluate “both” children and adults excluded. •The mean of 10mm was reported by 3 sites in each of those (i) reporting and (ii) not reporting false negatives.

